# Distinct profiles and antibody production by splenic and lymph node antibody-secreting cells in *Trypanosoma cruzi* infection

**DOI:** 10.64898/2026.10.24.761360

**Authors:** Laura Almada, Yamila Gazzoni, Juan Nahuel Quiroz, Julio C Gareca, Juan Mucci, Fernán Agüero, Eva V Acosta-Rodríguez, Carolina L Montes, Adriana Gruppi

**Affiliations:** Departamento de Bioquímica Clínica, Facultad de Ciencias Químicas (FCQ), Universidad Nacional de Córdoba, Córdoba, Argentina; Centro de Investigaciones en Bioquímica Clínica e Inmunología (CIBICI - CONICET), Córdoba, Argentina; Instituto de Investigaciones Biotecnológicas (IIBIO), Consejo Nacional de Investigaciones Científicas y Técnicas (CONICET), San Martín, Argentina; Escuela de Bio y Nanotecnología (EByN), Universidad Nacional de San Martín (UNSAM), San Martín, Argentina

**Keywords:** antibody secreting cells, tissue immunity, B cell response, *Trypanosoma cruzi*, Chagas disease

## Abstract

Evidence from bone marrow and intestinal lamina propria studies indicates that antibody-secreting cell (ASC) characteristics are shaped by the local microenvironment; yet whether secondary lymphoid organs harbor distinct ASC populations during infection remains unclear. Here, we investigated plasmablasts (PB) from the spleen and inguinal lymph nodes (LN) in acute *Trypanosoma cruzi* infection. Although PB from both tissues expressed canonical ASC markers, they displayed distinct transcriptional, phenotypic, and functional profiles. Splenic PB upregulated genes and proteins associated with inflammation, immunomodulation, and B-cell interactions. Activated and naïve B cells also exhibited tissue-specific transcriptional programs, suggesting that local cues influence B-cell differentiation. Consistent with these differences, spleens from infected mice contained more T follicular helper cells, whereas LN were enriched in regulatory T cells. High-dimensional flow cytometry identified multiple PB subsets differentially represented between tissues. Functionally, splenic PB secreted higher levels of IgM, IgA, and IgG, including both parasite-specific and autoreactive antibodies. In contrast, LN PB preferentially produced IgG against peptide 72 of the shed acute-phase antigen (SAPA), despite lower overall *T. cruzi*-specific antibody production. These findings demonstrate that secondary lymphoid organs harbor distinct PB populations adapted to their local microenvironments during acute infection.

## Introduction

Antibodies (Abs) play a central role in host defense against infections by recognizing and binding specific antigens (Ags) on pathogens, thereby limiting their spread, and promoting their elimination. The function of Abs is determined by the specialized characteristics of each isotype, as well as by the cells that produce them, which together shape the magnitude, localization, and quality of Abs-mediated immunity [1]–[4].

Antibody (Ab) isotype dominance and the characteristics of Ab-secreting cells (ASC), including plasmablasts (PB) and plasma cells (PC), are shaped by both Ag characteristics and the tissue microenvironment where the B-cell response is initiated [4],[5]. ASC found in secondary lymphoid organs (SLO), such as the spleen and lymph nodes (LN), more commonly produce IgG or IgM isotypes and are typically generated during systemic immune responses. Within the classical framework of B cell responses, following activation, B cells can adopt two major differentiation pathways: the extrafollicular response, in which activated B cells rapidly differentiate into short-lived PB that predominantly secrete low-affinity IgM, providing early humoral protection; alternatively, B cells may enter into the B cell follicle and generate germinal centers (GC), where they undergo somatic hypermutation and class-switch recombination, giving rise to high-affinity, class-switched ASC (e.g., IgG isotypes), as well as long-lived PC and memory B cells [6]. On the other hand, ASC present or originated in mucosal tissues produce large amounts of IgA and are shaped by continuous exposure to commensal microbiota and dietary Ags [7]. Notably, ASC localized in the intestine display distinctive features that reflect the unique microenvironment of the gut. These observations underscore the critical role of tissue-specific signals in imprinting distinct ASC phenotypes and Ab outputs.

ASC are not solely dedicated to immunoglobulin production but can also exert immunomodulatory functions [8]. It has been reported that subsets of ASC produce cytokines, such as IL-10 [9], IL-35 [10], IL-17 [11], and TGF-β [12], and/or surface-expressed proteins [13] or soluble mediators [14] that modulate T cell responses and inflammation. Therefore, it is possible that the environment not only shapes the characteristics of the Abs produced by ASC but also influences the expression of mediators responsible for executing regulatory functions [15].

It has become increasingly recognized in recent years that the environment of SLO actively orchestrate immune responses [16]–[19]. Interestingly, the highly ordered architecture of SLO is modified in response to infectious agents. In this sense, *Trypanosoma cruzi* infection is associated with profound changes in the composition, distribution, and relative proportions of hematopoietic immune cells in SLO [20],[21]. Acute *T. cruzi* infection is characterized by marked splenomegaly, lymphadenopathy, and extensive polyclonal B-cell activation, resulting in the production of autoreactive Abs alongside parasite-specific Abs (reviewed in [22]). In addition, the infection disrupts splenic architecture, leading to white pulp disorganization and GC collapse [23]. In the acute phase, a robust PB response emerges, reaching its peak before the peak of the GC response [13]. Furthermore, a population of T follicular cytotoxic cells, which is absent in non-infected mice, transiently arises during acute *T. cruzi* infection, modulating of early Ab responses [24].

We hypothesized that the distinct environments established in SLO during systemic infection with *T. cruzi* differentially shape ASC response, thereby influencing the magnitude and quality of the Abs. To test this hypothesis, we evaluated transcriptomic, phenotypic and functional characteristics of ASC present in anatomically distinct SLO, the spleen and inguinal LN, of *T. cruzi*–infected mice during the acute phase of infection.

## Results

### In the acute phase of infection, splenic and LN ASC from T. cruzi–infected mice show similar kinetic and are plasmablasts

To evaluate tissue-associated characteristics of ASC and considering that these cells are dynamic in terms of their generation and migration, we first analyzed the kinetics of ASC, together with cell populations potentially involved in their generation, such as GC B cells and follicular CD4 T (Tf) cells [25] in the spleen and inguinal LN of *T. cruzi*–infected mice. In this study, inguinal LN were the only LN analyzed; therefore, throughout the manuscript they are referred to simply as LN.

ASC were identified by flow cytometry using canonical markers (B220^interm^/^low^CD138^+^CD39^hi^CD98^+^, Figure 1A and Figure S1A) within the lymphocyte population. While ASC were rare in SLO of uninfected mice (0 dpi), during acute infection they increased with similar kinetics in spleen and LN, reaching a peak at 18– 20 dpi (Figure 1B). As previously reported with another *T. cruzi* strain such as the Y-Br strain [13], the ASC-increment preceded that of GC B cells identified as B220^+^CD138^neg^IgD^neg^Fas^+^Bcl-6^+^ (Figure 1C and Figure S1B), whose frequency peaked later (Figure 1D). The frequency of ASC declined by 28 dpi, whereas the frequency of GC B cells reached its maximum at 28 dpi and decreased thereafter. Once the acute phase of infection had resolved (60 and 150 dpi), ASC frequencies in both the spleen and LN were comparable to those observed in uninfected mice.

**Figure 1.**
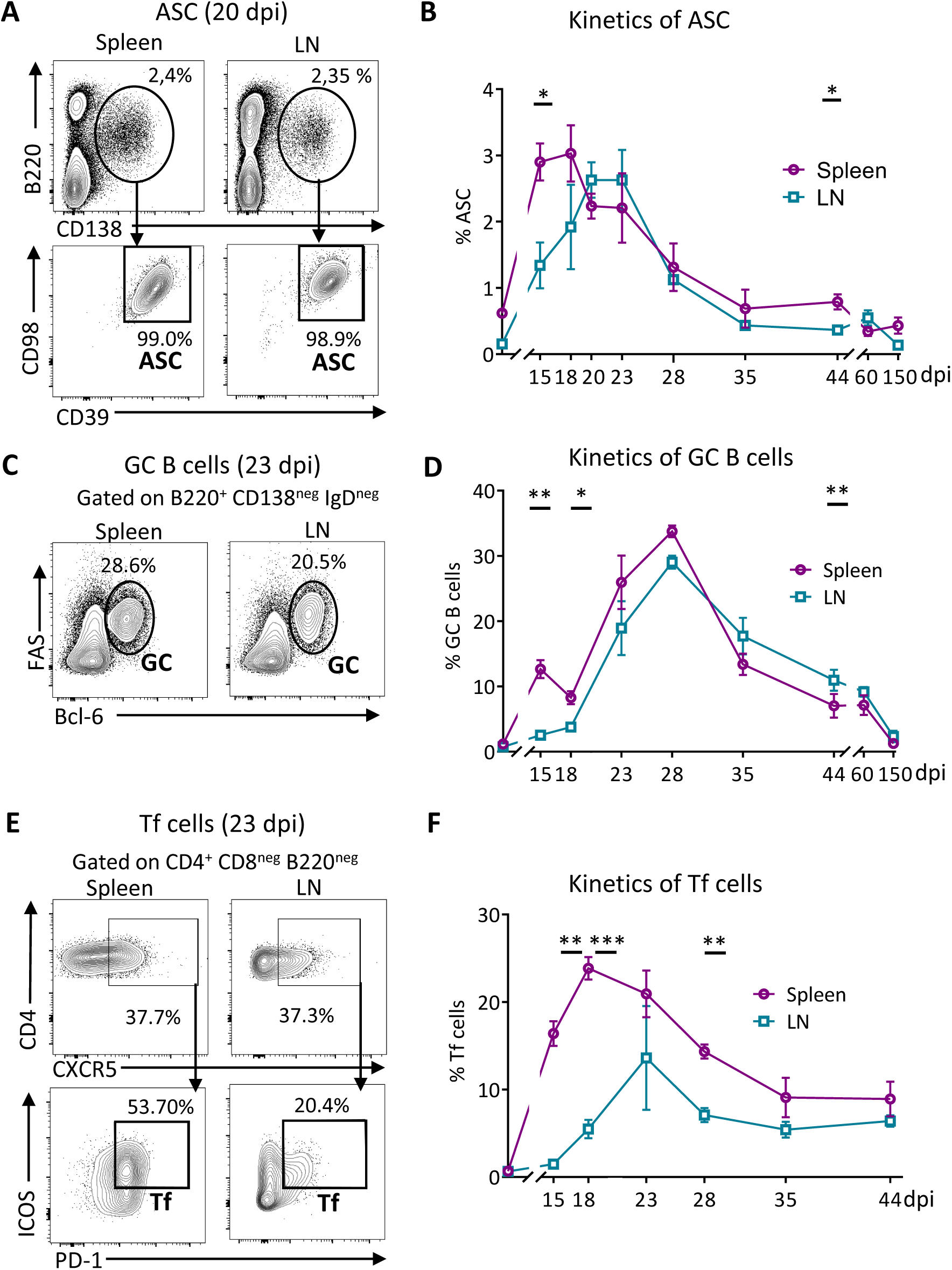
Kinetics of B cell response in spleen and LN of *T. cruzi*–infected mice. C57BL/6 mice were intraperitoneally injected with PBS (non-infected) or with 5,000 trypomastigotes of the *T. cruzi* Tulahuén strain (infected). Splenic and inguinal lymph node (LN) lymphocyte populations were analyzed by flow cytometry at different days post-infection (dpi). (A) Representative contour plots showing the percentage of ASC (B220^int/low^ CD138^+^ CD98^+^CD39^hi^) gated on live lymphocytes in spleen and LN at 20 dpi. (B) Kinetics of ASC frequency, expressed as a percentage of total lymphocytes, (C) Representative contour plots showing the percentage of germinal center (GC) B cells (B220 CD138 IgD Fas Bcl-6) gated on live lymphocytes in spleen and LN at 23 dpi. (D) Kinetics of GC B cell frequency, expressed as a percentage of B cells. (E) Representative contour plots showing the percentage of follicular CD4 T (Tf) cells (CD4 CXCR5 ICOS PD-1) gated on live lymphocytes in spleen and LN at 23 dpi. (F) Kinetics of Tf cell frequency, expressed as a percentage of CD4^+^ T cells. Kinetic of splenic and LN cells are indicated in violet and cyan, respectively. N ≥4 per group in each time point. Data are representative of 2 independent experiments. Statistical significance was determined by an unpaired t-test. (*p < 0.05**p < 0.01, ***p < 0.001)

For the evaluation of Tf cells, they were defined as CD4 CXCR5 ICOS PD-1 (Figure 1E and Figure S1C). Importantly, Tf cells were initially analyzed without discriminating based on Foxp3 expression; therefore, this population included both Foxp3^neg^ (T follicular helper - Tfh) and Foxp3^+^ (T follicular regulatory - Tfr) cells.

As shown in Figure 1F, Tf cells maximal frequency, within CD4 T cell population, occurred earlier and reached higher levels in the spleen than in LN. The peak frequency of splenic Tf cells occurred at 18 dpi and at 23 dpi in the LN, showing a temporal coincidence with the peak of ASC response in the spleen but not in the LN.

Early during infection (20 dpi), the majority of ASC corresponds to PB, as the entire population of ASC in both SLO expressed Ki-67 (Figure 2A). As infection progresses, the frequency of Ki-67^+^ASC decreased in both organs (Figure 2A, B). In Figure 2B, it can be observed that at 60 and 150 dpi, the frequencies of Ki-67-expressing cells were lower than during the acute phase of the infection, indicating that at these time points most ASC correspond to non-proliferating PC. Interestingly, the level of Ki-67 determined as mean fluorescence intensity (MFI) at 20 dpi, but not the frequency, was significantly reduced in ASC from LN in comparison to the splenic ASC (Figure 2C).

**Figure 2.**
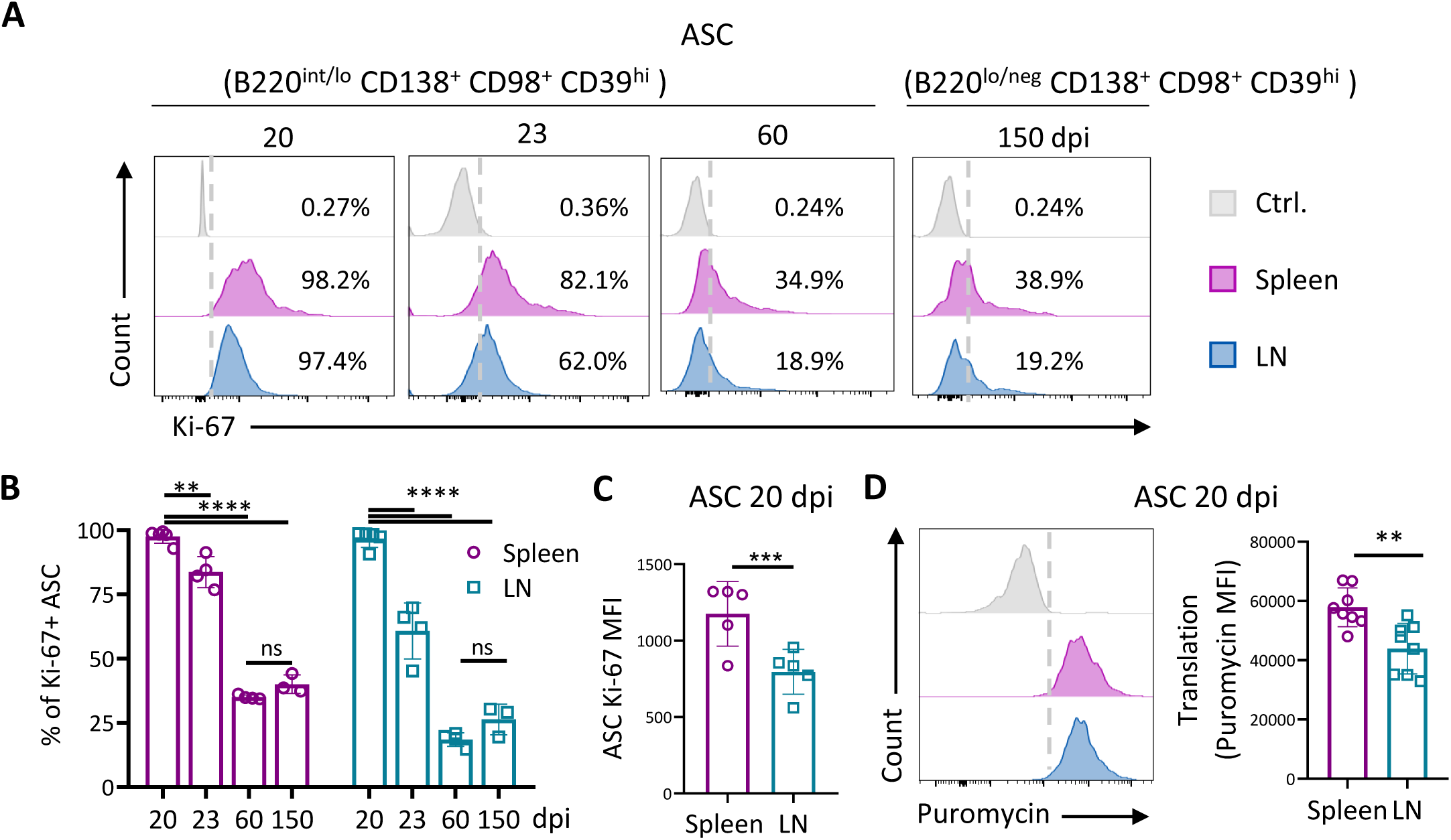
Expression of Ki-67 in ASC. Splenic and LN cells from *T. cruzi* infected mice were collected at different dpi (20, 23, 60 and 150) and the expression of Ki-67 in ASC was analyzed by flow cytometry. **(A-B)** Representative histograms and statistical analysis of the frequency of Ki-67^+^ASC (B220^int/low/neg^ CD138^+^ CD98^+^ CD39^hi^, gated on live lymphocytes) in spleen (violet) and LN (blue) of *T. cruzi* infected mice. **(C)** Expression of Ki-67, determined as mean fluorescence intensity (MFI), in ASC from the spleen and LN obtained at 20 dpi. **(D)** Representative histogram and statistical analysis of puromycin incorporation in ASC present in the spleen and LN obtained from infected mice at 20 dpi. N=8, performed in 2 independent experiments. **(A-B)** Data are presented as mean ± SD; N = ≥3 mice. Data are representative of 3 independent experiments. Statistical significance was determined by a paired t-test (**p < 0.01, ***p < 0.001, ****p < 0.0001). ns: no significant.

In addition, we observed that most ASC present in spleen and LN incorporated puromycin, reflecting active protein synthesis (Figure 2D). Splenic ASC exhibited higher puromycin incorporation compared to LN ASC, indicating increased protein synthesis activity by splenic ASC.

Because ASC from both tissues were predominantly Ki67^+^ at 20 dpi, indicating a large PB response, we selected this time point to compare tissue-associated ASC characteristics while minimizing time-dependent variability, as ASC may progressively differentiate into PC. Accordingly, the cells included in the comparative analysis were PB isolated from the spleen and LN of *T. cruzi*-infected mice obtained at 20 dpi.

### The spleen and LN of infected mice exhibit differences in cells and cytokines involved in PB-generation

Considering that the signals of tissue environment are relevant for immunity [26], we initially decided to evaluate the characteristic of cells with the potential to give rise to PB, as well as cytokines, present in the spleen and LN from infected mice. To evaluate whether each SLO harbors PB precursors with particular characteristics, we performed bulk RNA-seq on sorted naïve and activated B cells isolated from the spleen and LN of mice at 20 dpi (gating strategy shown in Figure S1B).

Gene expression analysis identified 850 genes that were differentially expressed between naïve B cell populations from both SLO (Figure 3A), and 684 genes that were differentially expressed between activated B cells from the spleen and LN (Figure 3B), indicating that B cell populations that directly generate PB differ between these tissues.

**Figure 3:**
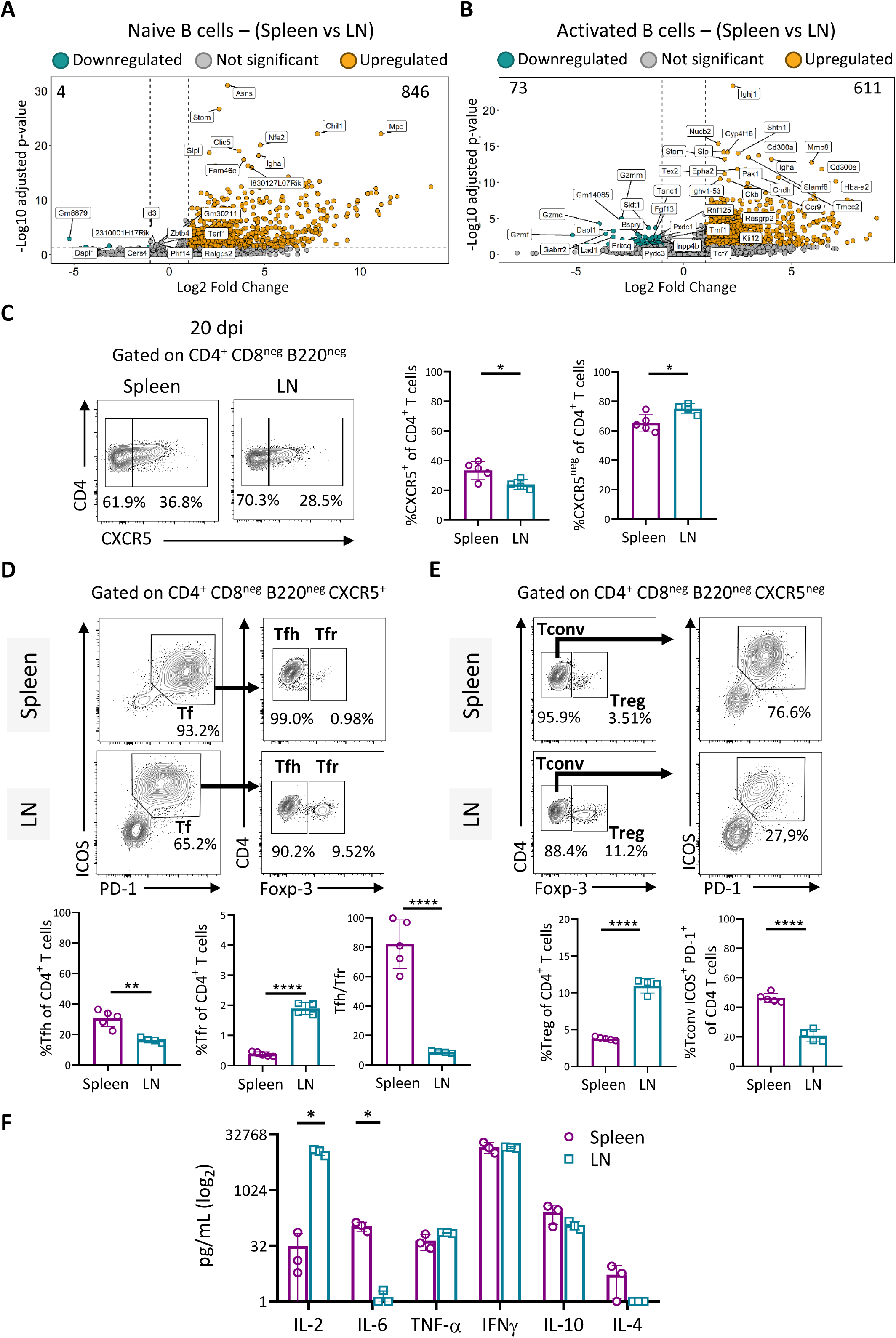
Cell populations involved in PB formation and cytokine production in the spleen and LN of *T. cruzi*-infected mice. Naïve and activated B cells were isolated by cell sorting from the spleen and LN from *T. cruzi*-infected mice at 20 dpi (three pools of 4-5 mice each, 3 biological replicates) and subjected to bulk RNA sequencing. Volcano plot displaying differentially expressed genes between **(A)** naïve B cells and **(B)** activated B cells from the spleen and LN. Significantly upregulated (orange dots) and downregulated (green dots) genes were defined by log2 fold change >1 and adjusted p-value < 0.05. Values in the upper corners indicate the number of differentially expressed genes. **(C)** Representative flow cytometry contour plots and statistical analysis of the frequency of CXCR5^+^ CD8^neg^B220^neg^ CD4^+^ T cells and CXCR5^neg^ CD8^neg^B220^neg^CD4^+^ T cells. **(D)** Representative flow cytometry contour plots and statistical analysis of the frequency of Tf (ICOS^+^PD-1^+^CXCR5^+^CD8^neg^B220^neg^ CD4^+^)Tfh (Foxp3^neg^ Tf) and Tfr cells (Foxp3^+^ Tf) from the spleen and LN of 20 dpi-infected mice. Tfh/Tfr ratio in CD4^+^ T cells from the spleen and LN. Each symbol represents an individual mouse. (E) Representative flow cytometry contour plots and statistical analysis of the frequency of Treg (Foxp3^+^CXCR5^neg^CD4^+^CD8^neg^B220^neg^) and ICOS^+^ PD-1^+^ Tconv (Tconv: Foxp-3^neg^ CD4^+^CD8^neg^B220^neg^) from the spleen and LN of 20 dpi-infected mice. Each symbol represents an individual mouse. (**E**) Statistical analysis of concentration (pg/ml) of IL-2, IL-6, TNFα, IFN-γ, IL-2, IL-10, and IL-4 in culture supernatants from single-cell suspensions of total spleen and LN cells after red blood cell lysis from 20 dpi infected mice, determined by ELISA. (**C-F)** Data is presented as mean ± SD; N = ≥3 mice. Data are representative of 3 independent experiments. Statistical significance was determined by a paired t-test (*p < 0.05, **p < 0.01, ****p < 0.0001). ns: no significant.

To assess the presence of T-cell subsets with helper or suppressive functions at the post-infection time point selected for the analysis of PB characteristics, we next performed a comprehensive phenotypic analysis of T-cell subsets. By flow cytometry, we evaluated the frequency of CXCR5^+^CD4^+^T cells and CXCR5^neg^ CD4^+^, which include conventional CD4^+^T and Treg cells. We observed, at 20 dpi, that spleen contained higher frequency of CXCR5^+^CD4^+^ T cells and as expected, lower frequency of CXCR5^neg^CD4^+^ T cells than LN (Figure 3C). Within the CXCR5^+^CD4^+^ T-cell population, Tf cells were identified based on ICOS and PD-1 expression and Tfh and Tfr were distinguished by Foxp3 expression. We observed that the SLO evaluated, at 20 dpi, displayed differences in Tfh and Tfr composition: LN exhibited a significantly lower frequency of Tfh cells, and a higher frequency of Tfr cells respect to the spleen (Figure 3D). Accordingly, the Tfh/Tfr ratio, which reflects the balance between helping vs restraining signals that influence Ab quality [27],[28], was significantly higher in the spleen than in the LN of infected mice (Figure 3D).

Analysis of the CXCR5^neg^CD4 T cell compartment (non-Tf cells) also revealed differences between tissues. In infected mice, Treg (CXCR5 Foxp3 CD4) cells were more frequent in the LN than in spleen, whereas CXCR5 Foxp3 ICOS PD-1 CD4 cells were more frequent in the spleen.

When assessing cytokine production by splenocytes and LN cells of infected mice, we detected significantly higher levels of IL-2 in LN-supernatants compared to the spleen-supernatants, whereas accordingly to high frequencies of Tfh cells, IL-6 levels were significantly increased in spleen-supernatants [29],[30]. The remaining cytokines analyzed, including TNF-α, IFN-γ, IL-10 and IL-4, showed comparable levels between both tissues (Figure 3F).

### Transcriptional Profiling Reveals Clear Tissue Segregation of Splenic and LN PB

To assess whether the cellular differences observed in the spleen and LN of infected mice are reflected in the PB populations of each tissue, we performed a transcriptional analysis of PB present in the different SLO. We conducted comparative bulk RNA-seq analysis of PB isolated by cell sorting (gating strategy Figure 1A) from the spleen and LN of 20 dpi–infected mice. In parallel, to simultaneously evaluate the contribution of tissue localization and B-cell differentiation stage to the transcriptional landscape, naïve and activated B cells from each organ were included in the principal component analysis (PCA). In Figure 4A, PCA plot shows the distribution of six different cell populations based on two principal components (PC1 and PC2). Naïve, activated B cells and PB segregated primarily according to differentiation state along PC1, which accounted for 82% of the total variance (Figure 4A). In contrast, PC2, which accounted for 10% of the total variance, separated the populations according to tissue origin. Importantly, while PB clustered separately from naïve and activated B cells due to their shared differentiation program, splenic and LN PB remained distinctly segregated along PC2, highlighting tissue-specific transcriptional differences. Hierarchical clustering further revealed a clear segregation of naïve and activated B cells from PB across the different SLO, with high intra-group similarity (Figure 4B).

**Figure 4:**
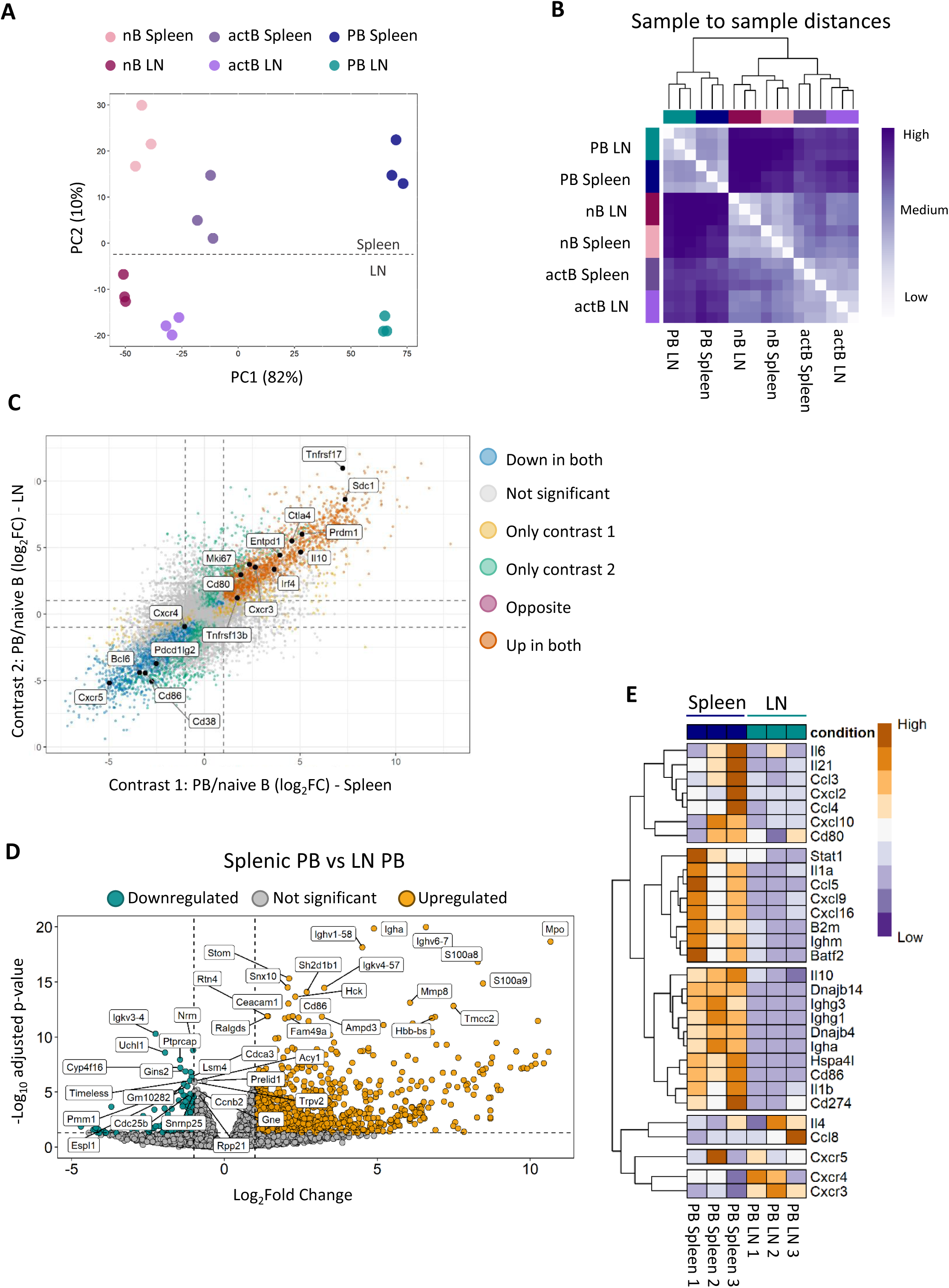
Whole-transcriptome profiling of naïve and activated B cells and PB across SLO. Naïve, activated B cells and PB were isolated by cell sorting from the spleen and LN from *T. cruzi*-infected mice at 20 dpi and subjected to bulk RNA sequencing (three pools of 4-5 mice each, 3 biological replicates). **(A)** PCA plot of DESeq2-normalized counts showing sample clustering by global gene expression. PC1 and PC2 explain 82% and 10% of the total variance, respectively. **(B)** Heatmap of sample-to-sample Euclidean distances with hierarchical clustering. Higher values indicate greater transcriptional dissimilarity, whereas lower values indicate greater similarity. **(C)** Four-way plot comparing log fold changes (PB vs naïve B cells) between spleen (contrast 1) and LN (contrast 2). Genes along the diagonal exhibit concordant regulation across tissues, whereas genes deviating from the diagonal display tissue-specific transcriptional responses. Selected genes of interest are labeled. **(D)** Volcano plot highlighting genes differentially expressed between PB from spleen and LN. The top 20 upregulated and downregulated genes are labeled. **(E)** Heatmap of normalized transcript counts for selected genes in PB isolated from spleen and LN. Genes with an adjusted p-value < 0.05 and an absolute log fold change > 1 were considered differentially expressed.

Consistent with these findings, the comparison of log fold changes of RNA expression in PB versus naïve B cells from spleen and LN revealed a shared transcriptional program associated with PB differentiation (Figure 4C). Canonical PB-associated genes, including Prdm1, Sdc1, Tnfrsf17, Irf4, Entpd1 and Ki67 were upregulated in PB from both tissues relative to naïve B cells, whereas genes associated with the naïve B-cell state, such as Bcl6 and Cxcr5 were downregulated. Although most hallmark PB genes were similarly enriched in splenic and LN PB, several transcripts displayed tissue-biased expression patterns, supporting the existence of tissue-specific transcriptional adaptations.

Focusing specifically on PB, we observed differential gene expression between PB from the spleen and LN. A total of 1065 transcripts were differentially expressed between the two PB populations, with 854 genes upregulated and 211 genes downregulated in splenic PB relative to LN-PB (Figure 4D).

Heatmap visualization of selected gene sets highlighted these differences, confirming distinct transcriptional programs in PB from both SLO (Figure 4E). PB from the spleen were enriched in transcripts associated with B cell cooperation and interaction such as IL-6 and IL21, including the transcription factor Batf, as well as costimulatory molecules such as CD80 and CD86 and effector molecules as immunoglobulins. Moreover, PB from the spleen showed increased expression of transcripts encoding inflammatory chemokines such as Ccl3, Ccl4, Ccl5, Cxcl9, and Cxcl16, together with elevated expression of transcripts encoding immunomodulatory molecules, including Il10, and Cd274.

Gene Ontology (GO) enrichment analysis revealed a significant overrepresentation of biological processes associated with immune functions in splenic PB compared with LN PB. Among the most enriched GO terms were *innate immune response*, *immune system process, inflammatory response,* phagocytosis *(engulfment), complement activation*, *chemotaxis, B-cell receptor signaling pathway, response to LPS, positive regulation of B cell activation, cellular response to interferon-beta* among others (Figure S2).

Collectively, these results indicate that although splenic and LN PB share a core transcriptional program associated with PB differentiation, each tissue harbors different PB population, with splenic PB displaying an enhanced inflammatory, immunomodulatory, and B cell–interactive profile compared with their LN counterparts.

### Splenic and LN PB comprise heterogeneous clusters with distinct levels of surface protein and transcription factors expression

To further characterize splenic and LN PB in the context of *T.cruzi* infection, we used multiparametric flow cytometry to assess the expression of various surface and intracellular proteins involved in diverse functions of ASC [8],[31]. We analyzed on PB (BB20^int^CD138^+^CD98^+^CD39^hi^) population the expression of co-stimulatory molecules involved in Ag-presentation (CD80 and CD86) [32]; chemokine receptors associated with migration and homing (CXCR3 and CXCR4) [33],[34]; and PD-L1, an immune checkpoint molecule involved in the regulation of immunity and inflammation [13]. We observed that both splenic and LN PB expressed these markers (Figure 5A); however, splenic PB exhibited significantly higher expression of these molecules, as determined by MFI, compared with LN PB (Figure 5A).

**Figure 5:**
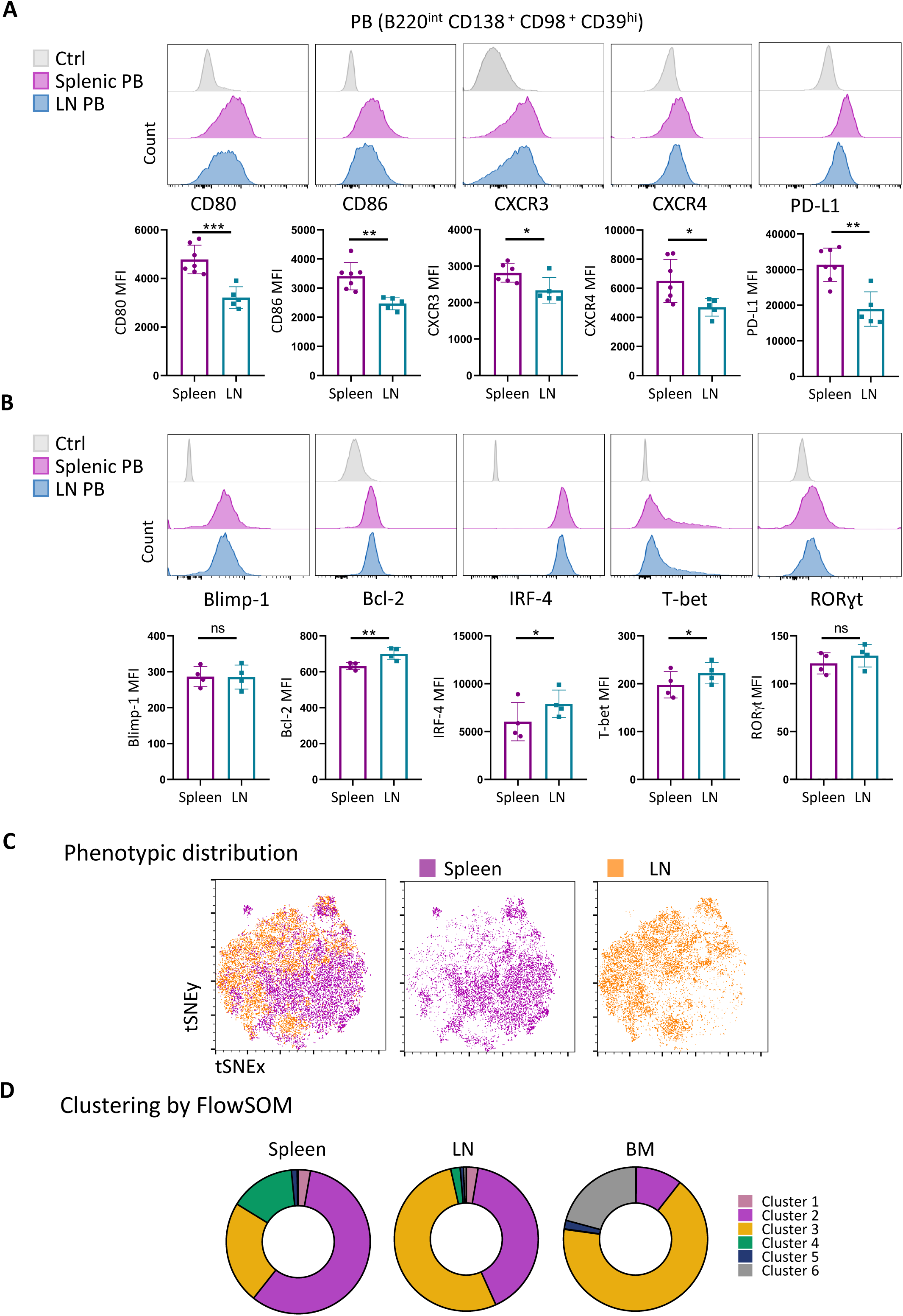
Phenotypic characterization of splenic and lymph node PB. **(A)** Representative histograms and statistical analysis of the MFI of CD80, CD86, CXCR3, CXCR4, and PD-L1 expression on splenic and lymph node PB (B220^int^CD138^+^CD98^+^CD39^hi^) from infected mice at 20 dpi. (**B)** Representative histograms and statistical analysis of the MFI of Blimp-1, Bcl-2, IRF-4, T-bet, and RORγt expression on splenic and lymph node PB from infected mice at 20 dpi. (**C)** t-SNE visualization showing the distribution of splenic and lymph node PB from infected mice at 20 dpi based on the expression of CD80, CD86, CXCR3, CXCR4, PD-L1, FAS, IgA, IgM, IgD, and Ki-67. Cells are colored according to tissue origin (spleen, violet; lymph node, orange. (**D)** FlowSOM analysis of splenic and lymph node PB from infected mice at 20 dpi, and bone marrow ASC (B220^lo/neg^CD138^int^Blimp-1^+^CD98^+^CD39^hi^) from infected mice at 130 dpi, based on the expression of MHC-I, CD80, CD86, FAS, PD-L1, CXCR3, and CXCR4. Graphs depict the percentage contribution of each cluster to the total PB or ASC population in the indicated tissues. (**A-B)** Data are presented as mean ± SD; N = ≥4 mice. Data are representative of 3 independent experiments. Statistical significance was determined by an unpaired t-test (*p < 0.05, **p < 0.01, ***p < 0.001). ns: no significant.

To determine whether PB from the different tissues expressed apoptosis-associated markers, we evaluated the expression of Fas and active caspase-3/9^+^ in ASC at 24 dpi, the time point at which the PB population begins to decline in each SLO. We observed that splenic ASC exhibited a higher frequency of Fas^+^ cells than LN PB (Figure S3A-B); however, this expression was not associated with increased apoptotic cell death, since the frequency of active caspase-3/9^+^ cells was similar between splenic and LN ASC (Figure S3B).

The analysis of the expression of transcription factors involved in the differentiation and maintenance of ASC, including Blimp-1 and IRF-4, as well as Bcl-2, a key survival factor that prevents programmed cell death, and T-bet and RORγt, revealed that PB from both SLO expressed these transcription factors. However, statistical analysis identified significant differences in the expression levels of Bcl-2, IRF-4, and T-bet (Figure 5B). Indeed, comparing to LN PB, splenic PB expressed higher levels of Bcl-2, IRF-4, and T-bet.

Next, we performed t-distributed Stochastic Neighbor Embedding (t-SNE) analysis to reduce the dimensionality of the high-parameter flow cytometry dataset and enable visualization of cellular heterogeneity. PB were defined as B220^lo/int^CD138^+^ CD98^+^CD39^hi^Blimp-1^+^ and by analyzing the expression of the markers CD80, CD86, CXCR3, CXCR4, PD-L1, FAS, IgA, IgD, IgM and Ki-67 together with MHC-I, the t-SNE plot shows a tissue-dependent segregation of PB, evidencing similarities among cells from the same tissue and differences across the two tissues (Figure 5C).

While our study focused on comparing splenic and LN PB at the acute phase of infection, bone-marrow ASC obtained at chronic phase were included in a multiparametric analysis as an additional reference population to further contextualize the phenotypic diversity identified within the ASC compartment. This approach enables a comparison of ASC phenotypes across tissues. For that, we performed a FlowSOM taking in account the surface expression of CD80, CD86, CXCR3, CXCR4, PD-L1, FAS, and MHC-I, within B220^lo/int/neg^CD138^+^CD98^+^CD39^hi^Blimp-1^+^ population. FlowSOM analysis of the integrated dataset (Figure 5D) identified six distinct ASC clusters that were differentially represented across tissues. The clusters were defined based on the expression of a combination of different levels of CD80, CD86, CXCR3, CXCR4, PD-L1, FAS, and MHC-I molecules. Splenic PB were mainly distributed between clusters 2 and 3, whereas chronic bone-marrow ASC were predominantly assigned to clusters 3 and 6. Notably, LN PB were largely represented by clusters 2 and 3, sharing major phenotypic clusters with both splenic PB and bone marrow ASC. Cluster 3 was highly represented across all ASC populations, while cluster 6 was almost exclusively associated with bone marrow. In contrast, clusters 1, 4, and 5 constituted only minor subsets. Together, these results reveal both shared and tissue-predominant ASC phenotypes across lymphoid organs.

### Splenic and LN PB differ in antibody isotype expression, secretion and specificity

To determine whether the phenotypic and transcriptional differences between splenic and LN PB are accompanied by differences in Ab production and isotype distribution, we first assessed the frequency of PB producing IgM, IgG e IgA. ELISpot assays, revealed a greater proportion of IgM- and IgA-secreting cells among splenic cells compared with their LN counterparts (Figure 6A), while LN cells displayed a higher frequency of IgG-secreting cells.

**Figure 6:**
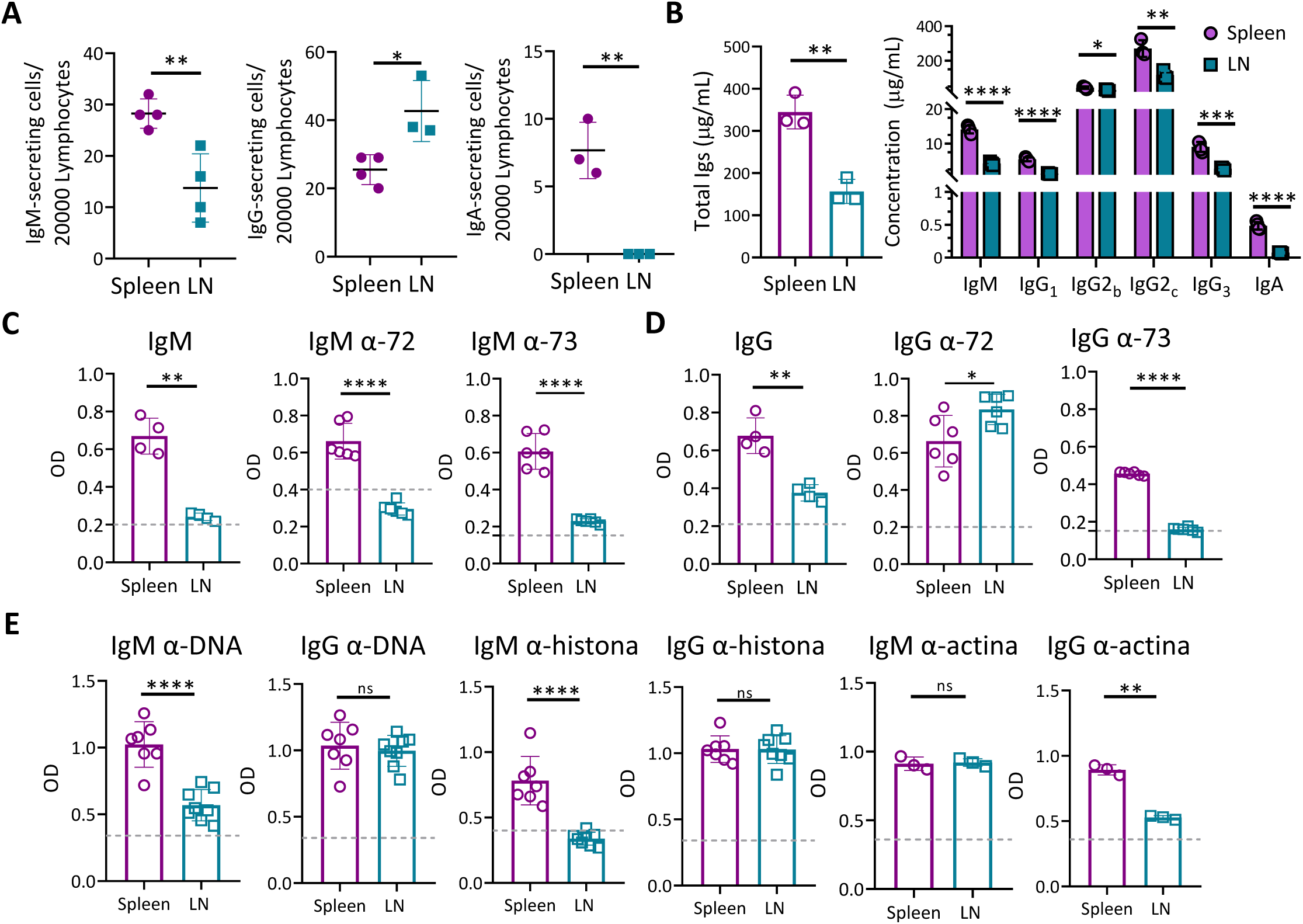
Antibody production by splenic and LN PB. **(A)** Numbers of IgM, IgG or IgA-secreting cells in 2 × 10^4^ cells from spleen or LN from 20 dpi-*T. cruzi* infected mice obtained by ELISpots. **(B-E)** 2 x10^5^ PB purified by cell sorting from spleen or LN of *T. cruzi* infected mice at 20 dpi were cultured per well in complete media plus BAFF (100 ng/mL) for 72 hours. (**B**) Bar plots show the total Abs levels in culture supernatants (left) and the distribution of individual Igs isotype (right) obtained by LEGENDplex. (**C-D)** Specific IgM and IgG anti-*T. cruzi*-in culture supernatants against trypmastigote homogenate measured by ELISA (left), and specific Abs for peptides 72 and 73 (middle and right, respectively) in the PB culture supernatants. (**E)** IgM and IgG autoantibodies against DNA, histone, and actin in PB culture supernatants were measured by ELISA. Data are representative of ≥2 independent experiments, N ≥ 3 per group per experiment. Statistical significance was determined by an unpaired t-test (*p < 0.05, **p < 0.01, ***p < 0.001, ****p < 0.0001. ns: no significant).

To assess the antibody-secreting capacity of each PB population, we next cultured same amount of sorted PB from the spleen and LN of *T cruzi* infected mice and measured the concentration of total Igs and IgM, IgA and IgG isotypes in the culture supernatants of PB cells (Figure 6B). We observed that the concentrations of IgM, IgG, and IgA isotypes were significantly lower in the supernatants of LN-PB than in those of splenic PB. Notably, IgA was not detected in the supernatants of LN-PB. These findings suggest that, despite the higher frequency of IgG-secreting PB in the LN, LN-derived PB exhibited a lower Ab secretory capacity than their splenic counterparts.

Next, we analyzed the specificity of Abs directed against parasite Ags. Parasite Ags included a total lysate of *T. cruzi* trypomastigotes and two parasite antigens, peptides 72 and 73, which we identified through a kinetic analysis as the main Ab targets present in the sera of mice during acute infection (Figure S4).

Reactivity against the longer SAPA peptide (here indicated as SAPA) was not evaluated in culture supernatants because Ab levels against these SAPA repeats were very low in sera from infected mice (Figure S4). We observed that culture supernatant from splenic PB exhibited higher levels of Abs specific for total *T. cruzi* Ags and antigenic peptides than LN PB culture supernatant, except for IgG anti-peptide 72 (Figure 6C and D). Despite the overall lower Ab production in LN PB, higher levels of anti-peptide 72 IgG were detected in these cultures (Figure 6D).

Because acute *T. cruzi* infection induces polyclonal B-cell activation and massive PB expansion, both of which have been associated with the generation of self-reactive antibodies [35],[36], we also assessed the presence of autoantibodies in PB culture supernatants. Splenic PB produced greater amounts of anti-DNA IgM, anti-histone IgM, and anti-actin IgG than LN PB. In contrast, the levels of anti-DNA IgG, anti-histone IgG, and anti-actin IgM were similar in culture supernatants from both tissues.

## Discussion

Here, we demonstrate that SLO harbor PB expressing canonical ASC markers while exhibiting particular features in each tissue. Splenic and LN PB were identified as heterogeneous populations that differed in their transcriptional profiles, phenotypic characteristics, and functional properties. Our findings indicate that in *T. cruzi* infection, PB, as well as activated and naïve B cells, exhibit distinct transcriptional characteristics in the spleen compared with the inguinal LN. Then, the distinctive features of PB populations in each tissue may be determined both by the cells that initially seeded the tissue, together with tissue-specific environmental factors generated during the course of infection.

Differences in Ag entry routes (blood-borne versus tissue-derived), local cytokines levels and stromal and immune cell composition are likely together responsible for shaping the magnitude, kinetics, and qualitative features of PB response. The differences observed in the cellular composition of the SLO analyzed here cannot be explained solely by their original stromal cell composition [37],[38], but also to changes of tissue architecture and cellular composition that arise as a consequence of *T. cruzi* infection [23],[24]. Our results reveal differences in the CD4 T-cell compartment between the spleen and LN during *T. cruzi* infection, suggesting that each SLO could provide distinct signals for B-cell and PB responses. Compared with LN, the spleen exhibited higher frequencies of Tfh cells and lower frequencies of Tfr cells, resulting in an increased Tfh/Tfr ratio which reflects the balance between helper and suppressive signals regulating Ab responses. In contrast, LN were enriched in non-Tfh and Treg cells. The results indicate that splenic environment could be more permissive for Ab production, including auto-reactive antibodies, whereas LN appear to maintain a more restrictive regulatory setting.

Importantly, these differences in CD4^+^ T cell populations could be associated with the distinct cytokine profiles detected in each tissue evaluated. LN cultures produced significantly higher levels of IL-2, whereas splenic cultures secreted higher amounts of IL-6. IL-2 is known to promote Treg cell survival and expansion while simultaneously inhibiting Tfh differentiation through STAT5-dependent pathways. Therefore, the elevated IL-2 levels detected in LN may contribute to the increased frequencies of Treg and Tfr cells and the reduced abundance of Tfh cells observed in this organ. Conversely, IL-6 is a key cytokine driving Tfh differentiation and maintenance by promoting Bcl-6 expression and supporting germinal center responses [39]. In the presence of elevated IL-2 levels, as we detected in LN, IL-6 signaling becomes necessary to sustain Tfh cell differentiation rather than Tfr development [29],[30]. The increased production of IL-6 in the spleen is therefore consistent with the higher frequencies of Tfh cells found in this tissue. Interestingly, despite the differences in IL-6 and IL-2 concentrations, the levels of TNF-α, IFN-γ, IL-10, and IL-4 were comparable between tissues, suggesting that the divergent responses observed in the spleen and LN are not primarily driven by differences in these cytokines.

PB from the spleen and LN exhibited distinct transcriptional profiles and patterns of surface molecule expression. Among these molecules, CXCR3 and CXCR4 are key chemokine receptors that direct ASC migration to specialized microenvironments. Although transcriptomic analysis did not reveal clear differences in Cxcr3 and Cxcr4 transcript levels in PB from both tissues, flow cytometry revealed significantly higher protein expression in splenic PB. Nevertheless, most PB in both tissues expressed these receptors, suggesting that each population retains the potential to migrate to either inflammatory sites or the bone marrow. Regarding molecules involved in Ag presentation and immune regulation, LN-derived PB expressed lower levels than splenic PB, indicating a lower activation state and, consequently, a reduced capacity for Ag-presentation and immunoregulatory functions.

FlowSOM analysis identified multiple ASC clusters in each tissue, further emphasizing the differences observed by transcriptomic and initial flow cytometry analysis. Notably, PB subpopulations distribution was more similar between LN and bone marrow than between spleen and bone marrow, suggesting that LN-derived PB more closely resemble the bone marrow PB compartment. This finding raises the possibility that LN-generated PB preferentially home to bone marrow niches that support long-term survival.

Our findings reinforce the concept that PB, as shown for PC [40], constitute heterogeneous populations and, additionally, provide new evidence that, across different SLO facing the same type of infection, the frequency of each subpopulation is differentially represented. We have identified several differences between PB from the spleen and LN of infected mice. The notion that a given cellular population can acquire distinct features in different lymphoid organs has been previously reported for ASC and other immune cell types. Accordingly, comparisons between PC from the small intestine lamina propria and those from the bone marrow have revealed the existence of tissue-specific gene transcription programs [41]. In line with this information, studies on innate lymphoid cells have established that immune cell phenotypes and functions can vary across SLO as a result of tissue-specific cues [42]. However, these comparisons were performed in tissues that are continuously exposed to microbial antigens and therefore remain under persistent antigenic stimulation. Here, we reported PB subpopulation characteristics in systemic tissues under the same infection.

The Ab secretion profile against *T. cruzi*- and self-antigens produced by splenic and LN-derived PB was distinct. In agreement with a microenvironment that favors Ab production because of higher frequency of Tfh cells, splenic PB exhibited a greater capacity for immunoglobulin secretion. Culture supernatants from splenic PB contained higher concentrations of IgM and IgA than those from LN-PB. IgA was nearly undetectable in the LN, suggesting that the inguinal LN microenvironment of *T. cruzi*-infected mice does not support the generation or persistence of IgA-producing cells. Despite exhibiting a higher frequency of IgG-secreting cells, determined by Elispot, LN-PB produced lower overall IgG levels in culture supernatants, suggesting a lower secretory capacity than splenic PB. IgG reactivity against peptide 72 was higher in LN PB supernatant than that observed in splenic PB supernatants, while similar levels was observed for anti-DNA and anti-histone IgG. These findings indicate that a reduction in total IgG production by LN PB does not necessarily result in diminished antigen-specific or autoreactive IgG responses. Although inguinal LN provide a more controlled immune environment, characterized by a higher frequency of Tregs, PB from this organ, similar to splenic PB, have the capacity to produce autoantibodies. Finally, an interesting observation is that PB from spleen and LN were capable of producing IgM but also substantial amounts of IgG of different isotypes, highlighting that these PB have undergone class switch recombination in both SLO.

In summary, our findings show that cell populations defined by conventional markers display site-specific phenotypic and functional heterogeneity, reflecting adaptation to their local microenvironment. These observations highlight that analyses restricted to a single SLO, as spleen or LN, are insufficient to infer the nature of systemic immune responses. The tissue differences, simultaneously with the naïve B cells that seed each tissue could contribute to the heterogeneous generation of PB response observed in *T. cruzi* infection and could influence the parasite-specific and autoreactive Ab responses. One possible interpretation is that functionally interconnected tissues may cooperate through the generation of distinct PB populations, leading to Ab responses with diverse isotypes and specificities thereby contributing to the early control of the pathogen. Our work highlight PB compartmentalize responses, likely allowing B cell-mediated immunity to be appropriately adapted in both magnitude and quality to each anatomical site according to the infection.

## Material and methods

### Mice and experimental Infection

All animal procedures were evaluated, approved, and performed in accordance with the guidelines of the Institutional Committee for the Care and Use of Laboratory Animals at the “Facultad de Ciencias Químicas”, Universidad Nacional de Córdoba (Approval No. HCD RD-2021-2033-E-UNC-DEC#FCQ). The C57BL/6 mouse colony, originally obtained from the “Facultad de Veterinaria”, Universidad Nacional de La Plata (La Plata, Argentina), has been bred and maintained at the CIBICI-CONICET, FCQ-UNC Animal Facility for more than seven years.

Both male and female age-matched mice (8–12 weeks old) were utilized and kept under a 12 h light/dark cycle with controlled temperature and humidity. Animals were housed under specific pathogen-free conditions with unrestricted access to food and water. For each experiment, mice were placed in cages on the same rack and at comparable heights. Animal facility personnel were blinded to group allocation. In addition, prior to tissue collection, an independent individual assigned identification codes to each mouse to ensure blinded sample processing and data analysis. Mice exhibiting signs of severe illness, distress, or marked weight loss were humanely euthanized.

For experimental infection, mice were inoculated intraperitoneally with 5 × 10³ *T. cruzi* (Tulahuén strain) trypomastigotes in 0.2 ml PBS [43]. Control littermates received PBS alone and were processed in parallel.

### Cell Preparation

Spleen and inguinal LN were harvested at different days post-infection (dpi) for immunological analyses. Bone marrow cells from infected mice were obtained at 150 dpi by flushing femurs and tibias with RPMI-1640 medium. Tissues were mechanically dissociated by passage through a 70-µm cell strainer. Red blood cells were removed by incubation for 5 min in Tris–ammonium chloride lysis buffer. Live cell counts were assessed by trypan blue exclusion using a hemocytometer.

### Flow cytometry

For surface staining, single-cell suspensions were washed first in ice-cold PBS and incubated for ten minutes with a Live/Dead Staining. Cells were washed in PBS and then incubated with fluorochrome-labeled Abs for 30 min at 4°C (surface staining). For intracellular staining cells were fixed and permeabilized with BD Cytofix/Cytoperm and Perm/Wash (BD Biosciences) according to the manufacturer’s instructions. Data were collected on a BD FACSCanto II, BD Fortessa X20 and Northern Lights (Cytek) and were analyzed using the FlowJo software (TreeStar). The specific cell populations and gating strategy in each case are described in Figures and supplementary Figures (appropriately mentioned in the corresponding Result section). High-dimensional flow cytometry data were analyzed using t-distributed stochastic neighbor embedding (t-SNE) for dimensionality reduction. Cell clustering was performed using the FlowSOM algorithm based on the expression of the indicated markers. The resulting clusters were visualized and characterized with ClusterExplorer, allowing the assessment of marker expression patterns and the identification of phenotypically distinct cell populations. Flow cytometry data were acquired, analyzed, and reported in accordance with the EJI Guidelines for the use of flow cytometry and cell sorting in immunological studies.

### Antibodies

The following anti-mouse antibodies were used for flow cytometry: CD19-PE (6D5), CD19-BV605 (6D5), CD3-AlexaFluor700 (17A2), CD3-APCCY7 (17A2), CD4-APC-Cy7 (GK1.5), CD4-Pacific Blue(RM4-4), CD8-FITC (5H10-1), Streptavidin BV605, PD-1-BV421 (29F.1A12), PD-L1-PECy7 (10F.9G2), Bcl6-PE-Dazzle594 (7D1), ICOS-AlexaFluor488 (C398.4A), B220-APCCy7 (RA3-6B2), B220 PE(RA3-6B2), B220-PECy5(RA3-6B2), CD38-Pacific Blue(90), IgD-APCCy7(11-26c.2a), IgD-BV650(11-26c.2a), CD98-PE(RL388), CD98-PerCPCy5.5(RL388), CD98-PECy7(RL388), CD86-BV785(GL-1), CXCR3-BV510(173), CXCR4-BV605(L276F12), IgM-BV711(RMM-1) and Live Dead Aqua 430 and Zombie Aqua 510 were from Biolegend;, IgD-FITC (11-26c), CD8-PECy7 (53-6.7), PD-L1-PE, (MIH5), CD3-APC(145-2C11), Streptavidin-PE, Foxp3-FITC (FJK-16s), Ki-67-PE(SolA15), CD39-PE-Cy7(24DMS1), CD39-AF700(24DMS1), CD80-PerCPeF710(16-10A1), IRF-4-PerCp-eFluor710(3E4), T-bet-PECy7(4B10), IgA-FITC(mA-6E1) and MHC-I SB436(AF6-88.5.5.3) were from eBioscience; CD138-BV605 (281-2), CD138-APC (281-2), CD138-PE (281-2), CD38-BV421 (90/CD38), CXCR5-Biotin (2G8), Ki-67-AlexaFluor647 (B56), Blimp-1-PE-CF594 (5E7), B220-BV711(RA3-6B2), IgD-FITC(11.26c.2a), FAS-PECF594(JO2), FAS-PECy7(JO2), Bcl-6AF647(K112-91), Bcl-6-PECF594(K112-91), PD-L1-BV421(MIH5), Bcl-2-FITC(10C4) and RORγt-PE(Q31-378), FAS-PECy7(Jo2) were from BD Biosciences.

### Puromycin Assay

One x10^6^ cells isolated from the spleen or LN from infected mice obtained at 20 dpi were cultured in RPMI medium with or without puromycin (10 μg/mL). After incubation for 20 min at 37°C, cells were washed, permeabilized as previously described and stained with an Alexa Fluor 647-conjugated anti-puromycin antibody for flow cytometry detection.

### Cell Sorting

Spleen and LN from *T. cruzi*-infected mice were collected at 20 dpi. From live lymphocytes, PB (B220^int^CD138^+^, all ASC in these tissues at 20 dpi are PB), naïve B cells (B220^+^IgD^+^), and activated B cells (B220^+^ IgD^neg^CD138^neg^), were sorted using a BD FACSAria II cell sorter and used for RNA sequencing. For this purpose, three pools were generated per group for both spleen and lymph nodes, with each pool comprising cells isolated from the same 4–6 animals (n = 3 per tissue, per group)

For PB culture experiments, PB were further defined as B220^int^CD138^+^CD39^hi^CD98^+^ and sorted from the same samples using the same instrument.

### RNA sequencing

Total RNA was extracted using the Arcturus® PicoPure® RNA Isolation Kit according to the manufacturer’s instructions from PB, naïve and activated B cells from three *T. cruzi*-infected mice sorted from inguinal LN and spleen at 20 dpi, representing three biological replicates for each sample type, all obtained at the same infection time. RNA sequencing (RNA-seq) was carried out by GENEWIZ Multiomics & Synthesis Solutions (Azenta Life Sciences, NJ, USA). Poly(A) selection was used to remove rRNA, and libraries were prepared using an ultra-low input protocol by GENEWIZ. For RNA-seq analysis, sequencing reads were trimmed to remove adapter sequences and low-quality bases using Trimmomatic (v0.36). Trimmed reads were aligned to the Mus musculus reference genome (GRCm38, Ensembl release) using STAR (v2.5.2b), and gene-level raw count matrices were subsequently generated. Genes with low read counts (<10 per group) were removed prior to DESeq2 (v1.42.1) differential expression modelling, with cell type and tissue included as covariates in the design formula. Statistical significance and log2 fold changes were estimated using the Wald test [44]. Genes with an adjusted p-value < 0.05 and an absolute log2 fold change > 1 were considered differentially expressed in each comparison. Euclidean distances were calculated based on the variance-stabilized gene expression counts. Data visualization and graphical representations were generated in RStudio (v2023.06.0).

### Cell cultures

Two x10^6^ splenocytes or cells from LN from were cultured with complete media plus PMA-ionomycin for 5 h. Culture supernatants were collected for Cytokine quantification. Two x10^5^ ASC purified by cell sorting were cultured with complete media plus BAFF (100 ng/mL) for 72 h. Culture supernatants were collected for Ab-identification and quantification.

### Cytokine quantification

Cytokines (IL-2, IL-6, TNF-α, IFNγ, IL-10 and IL-4) were quantified using a bead-base multiplex assay by FACS (LEGENDplexTM, Mouse Cytokine Inflammation Panel) in the culture supernatant of 2 x10^6^ cells from splenocytes or LN cells obtained at 20 dpi. Supernatants and standards were run in duplicates in plates according to the manufacturer’s instructions (BioLegend). The samples were transferred from plate to tube and read using a FACSCanto II cytometer and data were analyzed with LEGENDplexTM Data Analysis Software.

### Immunoglobulin concentrations

Immunoglobulins (IgM, IgG1, IgG2b, IgG2c, IgG3, and IgA) were quantified using a bead-base multiplex assay by FACS (LEGENDplexTM), Mouse Immunoglobulin Isotyping Panel, 6 plex) in the culture supernatant of ASC. Supernatants and standards were run in duplicates in plates according to the manufacturer’s instructions (BioLegend). The samples were transferred from plate to tube and read using a FACSCanto II cytometer and data were analyzed with LEGENDplexTM Data Analysis Software.

### Determination of *T. cruzi* specific Abs

*T. cruzi*-specific Abs were quantified by ELISA using *T. cruzi* lysate-sensitized plates from Wiener lab (Rosario, Argentina). The wells were incubated with supernatants from PB sorted from spleen or LN, for 2 h at room temperature (RT). Peroxidase-conjugated anti-mouse IgM or IgG isotypes were added and incubated for 1:30 h at room temperature. The reaction was developed with TMB Substrate Reagent (BD Biosciences) and read in a Microplate Reader 450 from Bio-Rad. The ELISA test was considered positive if the mean OD value was two standard deviations above the mean value for the control supernatant assayed in parallel.

Abs against *T. cruzi* peptides were quantified by ELISA using synthetic peptides to sensitize the plates. Synthetic peptides 72 and 73 were produced by Schafer-N (Copenhagen, Denmark), and the synthetic peptide named SAPA was synthesized by JPT Peptide Technologies GmbH (Berlin, Germany). Peptides 72 (sequence TPADSSAHSTPSTPAC) and 73 (sequence EPKSAEPKPAEPKSAC) correspond to peptides pc4 and pc3 respectively in Mucci et al. 2017 [45]. The SAPA peptide is a 28mer (sequence SSSDSSAHGTPSTPVDSTAHGTPSTPAC) spanning two repeats of the SAPA antigen. All peptides were purified by HPLC (>90% purity), characterized by mass spectroscopy, and coupled to maleimide-activated BSA as described [45]. The wells were blocked and incubated with supernatants from PB from spleen or LN or mouse serum, for 2 h at room temperature. Peroxidase-conjugated anti-mouse IgM or IgG isotypes were added and incubated for 1:30 h at room temperature. The reaction was developed with TMB Substrate Reagent (BD Biosciences) and read in a Microplate Reader 450 from Bio-Rad. In the case of peptide-specific IgG from serum, an anti-mouse IgG conjugated with FITC was used and the fluorescence was read in a fluororeader at 488nm.

### Determination of auto-Abs

To evaluate specific auto-Abs, 96-well plates were coated with 5 μg/mL of calf thymus double-stranded DNA (D3664-5×2MG, Sigma-Aldrich), 5 μg/mL of histones (ATH01-02, Arotec Diagnostic Limited), or 5 μg/mL of actin (A0398, Sigma-Aldrich), diluted in carbonate buffer (pH 9.5), and incubated overnight at 4 °C. Then, plates were washed three times with PBS containing 0.05% Tween-20 (PBS-T) and blocked with PBS containing 3% BSA for 2 h at 37 °C. Culture supernatants were diluted 1:2 in PBS containing 1,5% BSA and incubated overnight at 4 °C. Peroxidase-conjugated anti-mouse IgM or IgG isotypes were added and incubated for 1:30 h at room temperature. The reaction was developed with TMB Substrate Reagent (BD Biosciences) and read in a Microplate Reader 450 from Bio-Rad. The ELISA test was considered positive if the mean OD value was higher than twice the mean value for the control assayed in parallel. Results were expressed as absorbance values (450 nm).

### ELISpot assay

To measure the frequency of ASC, ELISpot multiscreen filtration plates (Millipore) were activated with absolute ethanol, washed with PBS, and coated overnight at 4°C with 10 μg of the isotype-specific goat anti-mouse IgM, IgG and IgA (Southern Biotech), for total ASC detection, overnight at 4°C and blocked with 1% BSA. Plates were subsequently blocked for 2 h with complete medium and incubated for 24 h at 37°C with serial dilutions of single-cell suspensions. Plates were washed with PBS 0.01% Tween-20 and incubated for 1h with 1mg/ml of biotin-conjugated anti-IgM, IgG or IgA (SouthernBiotech) diluted in PBS 1%BSA. Then, plates were washed with 0.1% Tween 20 and incubated with 1 mg/ml ExtrAvidin-alkaline phosphatase (Sigma-Aldrich). Spots were detected using BCIP/NBT (Sigma-Aldrich).

### Statistics

Statistical significance through comparison of mean values was assessed by a two-tailed Student’s t-test, one-way or two-way ANOVA followed by Bonferroni’s posttest using GraphPad-Prism software.

## Data availability

The bulk RNA-seq data have been deposited in the NCBI BioProject database under accession number PRJNA1345747.

## ACKNOWLEDGEMENTS

We thank the staff from Cell Sorter and Cytometry Core and Animal facility from CIBICI-CONICET. This work was supported by grants from the Agencia Nacional de Promoción Científica y Técnica (PICT 2018-01494), SECYT-UNC (33620190100020CB), CONICET (PIP 201511220150100560CO), and R01AI116432 from NIH. EAR, CLM, and AG are Researchers from CONICET. LA, YG and JCG thank CONICET for the fellowship awarded.

