## Supplemental Fig. 1 for "Distinct profiles and antibody production by splenic and lymph node antibody-secreting cells in *Trypanosoma cruzi* infection"

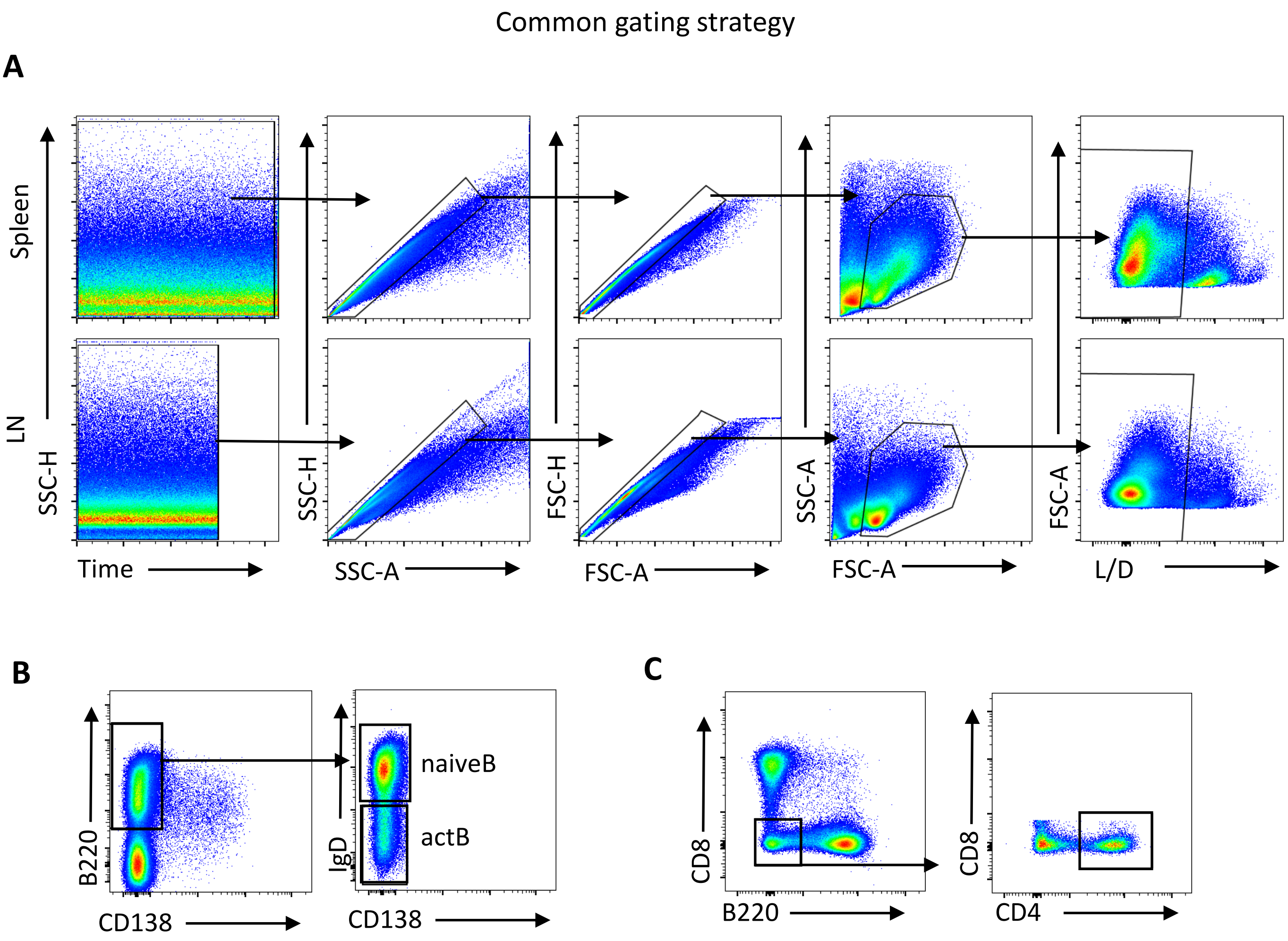


**Supplementary Figure 1. Flow cytometry gating strategies.**

**(A)** Flow cytometry gating strategy used for the selection of singlets, lymphocytes, and live cells (L/D). **(B)** Gating strategy used for the identification and sorting of naïve B cells and activated B cells (actB). (**C)** Gating strategy used for the identification of CD4 T cells.
