## Supplemental Fig. 2 for "Distinct profiles and antibody production by splenic and lymph node antibody-secreting cells in *Trypanosoma cruzi* infection"

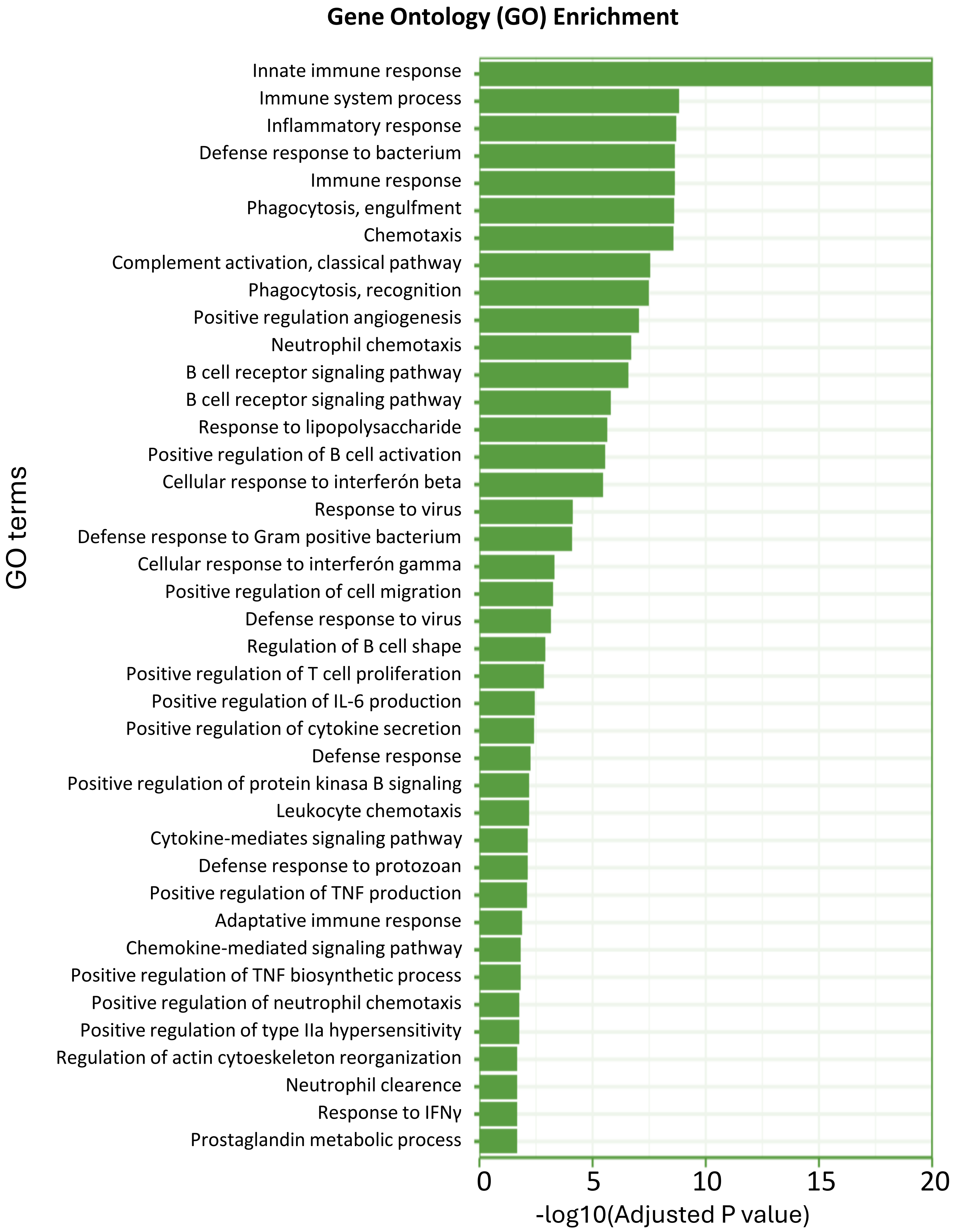


**Supplementary Figure 2. Gene ontology analysis.** Significantly differentially expressed genes were clustered by their gene ontology and the enrichment of gene ontology terms was tested using Fisher exact test (GeneSCF v1.1-p2). Figure shows gene ontology terms that are significantly enriched in splenic PB compared with LN PB, with an adjusted P-value less than 0.05 in the differentially expressed gene sets.
