## Supplemental Fig. 3 for "Distinct profiles and antibody production by splenic and lymph node antibody-secreting cells in *Trypanosoma cruzi* infection"

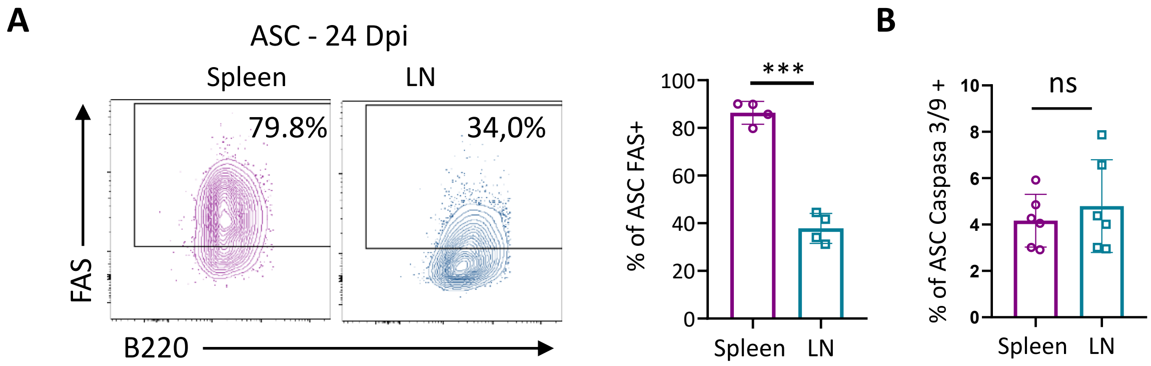


**Supplementary Figure 3. Expression of FAS and Caspasa 3/9 by ASC**
**(A)** Representative contour plots and statistical analysis of Fas expression on splenic and lymph node ASC from infected mice at 24 dpi. (B) Bar graph showing the percentage of splenic and lymph node ASCs expressing caspase-3/9 at 24 dpi. Data are representative of 3 independent experiments, N ≥ 4 per group per experiment. Statistical significance was determined by a paired t-test (***$p < 0.001$, ns: no significant).
