## Supplemental Fig. 4 for "Distinct profiles and antibody production by splenic and lymph node antibody-secreting cells in *Trypanosoma cruzi* infection"

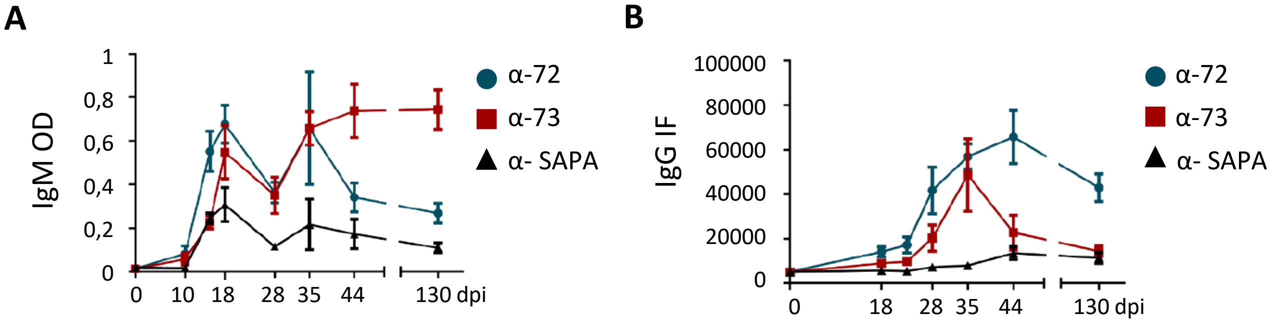


**Supplementary Figure 4. Kinetics of parasite-specific antibody responses against *T. cruzi* antigens.**

**(A)** Kinetics of parasite-specific IgM antibodies recognizing peptides 72, 73, and SAPA in plasma samples from *T. cruzi*-infected mice, assessed by ELISA. **(B)** Kinetics of parasite-specific IgG antibodies recognizing peptides 72, 73, and SAPA in plasma samples from *T. cruzi*-infected mice, assessed by a fluorescence-based assay.
